# Dynamics of thalamic directional coding under vestibular imbalance

**DOI:** 10.1101/2025.03.31.646416

**Authors:** Nada El Mahmoudi, Antoine Huret, Célia Laurent, Jean Laurens, Pierre-Yves Jacob, Francesca Sargolini

**Affiliations:** Aix-Marseille Université, CNRS, CRPN (Centre de Recherche en Psychologie et Neurosciences) – UMR 7077, Marseille, 13003 France; Ernst Strüngmann Institute (ESI) for Neuroscience in Cooperation with Max Planck Society, Frankfurt am Main, 60528, Germany

## Abstract

Head direction cells (HDCs) encode the animal’s orientation in space and form a core component of the brain’s navigation system. While their dependence on vestibular input is well established, how directional circuits respond to unilateral vestibular loss (UVL) — the most frequent and ecologically relevant form of vestibular imbalance — remains largely unknown. UVL offers a powerful model to investigate how spatial circuits adapt to asymmetric sensory disruption and partial deafferentation. To explore this, we examined the impact of UVL on anterior thalamic nuclei (ATN) activity and spatially tuned neurons in freely moving rats following unilateral vestibular neurectomy (UVN). UVN induced long-lasting alterations in ATN firing dynamics, including reduced theta modulation, diminished burst firing, and selective disruption across functionally defined neuronal classes: head-direction and speed-modulated cells were strongly affected, while angular head velocity and position cells remained largely preserved. Despite initial degradation, HDCs persisted and progressively regained directional tuning. Crucially, spike waveform analysis revealed two distinct HDC subtypes with markedly different vulnerabilities: one subtype showed reduced prevalence and degraded tuning, whereas the other remained resilient and supported the recovery of directional coding. These findings uncover a previously unrecognized heterogeneity within the head direction system and show that compensation following UVL is partial, cell type–specific, and functionally selective. Together, they offer new insight into sensory plasticity within thalamic navigation circuits and provide a framework to understand spatial deficits associated with vestibular imbalance.

## Introduction

Our ability to navigate the world relies on a network of spatially tuned neurons, among which head direction cells (HDCs) play a crucial role by encoding the animal’s direction in space. These neurons, located in a distributed network encompassing the lateral mammillary bodies, anterior thalamic nuclei (ATN), retrosplenial cortex, and presubiculum, are thought to provide the neural basis of our internal compass (Cullen and Taube, 2017). Crucially, HDCs depend on vestibular input to generate their directional signal, making them particularly vulnerable to disruptions in vestibular processing (Calton and Taube, 2005; Muir et al., 2009; Valerio and Taube, 2016). Given this strong vestibular dependence, HDCs offer a powerful model system to investigate how directional representations are impacted by vestibular dysfunction. While the effects of bilateral vestibular lesions on head direction coding have been studied (Stackman and Taube, 1998; Valerio and Taube, 2016; Yoder and Taube, 2009), the impact of unilateral vestibular loss (UVL) on this system remains largely unexplored. Yet, UVL provides a particularly informative model to investigate how directional circuits respond to asymmetric sensory disruption — a condition that introduces an imbalance between hemispheric inputs rather than complete bilateral deprivation. Moreover, UVL represents the most common form of vestibular dysfunction in humans (Brandt and Dieterich, 2017), further highlighting the importance of understanding how navigation circuits adapt to such perturbations.

UVL induces a characteristic vestibular syndrome involving posturo-locomotor, oculomotor, vegetative, and perceptual-cognitive deficits (Bronstein and Dieterich, 2019; Curthoys and Halmagyi, 1995; Uffer and Hegemann, 2016). These symptoms originate from an acute electrophysiological imbalance between the ipsi- and contralesional vestibular nuclei (VNs) (Dutia, 2010; Mccabe and Ryu, 1969; Precht et al., 1966), which progressively resolves over time. This recovery process, known as vestibular compensation, involves dynamic neuroplastic changes at multiple levels of the central nervous system and ultimately supports the restoration of electrophysiological balance and functional recovery (Beraneck and Idoux, 2012; Lacour et al., 2016).

While sensorimotor functions such as balance and posture typically recover, spatial cognitive deficits often persist (Chapuis et al., 1992; Gammeri et al., 2022; Guidetti et al., 2020, 2007), suggesting that compensation is incomplete at higher integrative levels. In a recent study, we showed that UVL induces long-lasting impairments in both spatial working and reference memory, associated with deficits in hippocampal plasticity (El Mahmoudi et al., 2023). These findings highlight the limits of compensation in cognitive domains and underscore the need to investigate how navigation circuits, particularly the head direction system, adapt — or fail to adapt —after unilateral vestibular loss.

Within the head direction circuit, the ATN serves as a critical relay, receiving vestibular inputs via a subcortical pathway including the medial vestibular nucleus (MVN), dorsal tegmental nucleus, and lateral mammillary nuclei (Cullen and Taube, 2017; Mehlman et al., 2021). Lesions of the ATN disrupt both directional coding and spatial memory (Aggleton and Nelson, 2015) making this region an ideal target to investigate the impact of UVL on the directional signal.

In this study, we sought to characterize how UVL and subsequent vestibular compensation affect neuronal activity within the ATN, with a particular focus on head direction cells. Using a rat model of unilateral vestibular neurectomy (UVN), we tracked the temporal dynamics of both behavioral and electrophysiological responses to assess the overall impact of UVL on ATN activity, and more specifically, on directional coding.

## Results

### UVN induces behavioral and *c-fos* changes in the ipsilesional ATN

Because MVN–ATN projections are bilateral, we first assessed whether the ATN shows lateralized activation following UVN. To this end, we quantified the number of *c-fos* immunoreactive cells in the MVN and the ATN, 24 hours after UVN, in both sham and lesioned rats. This analysis aimed to determine whether the lesion triggered ATN activation, and whether this response was lateralized. UVN triggered a robust increase in *c-fos* expression in both structures (**Fig.1B**, MVN: Kruskal-Wallis (KW) statistic = 21.14, p < 0.001; ATN: KW statistic = 22.6, p < 0.001), with a particularly marked response on the ipsilesional side (Dunn’s multiple comparisons test: sham ipsi vs UVN ipsi, MVN adj-p < 0.001; ATN adj-p < 0.05). This ipsilateral predominance guided our decision to perform subsequent electrophysiological recordings in the ipsilesional ATN.

**Figure 1:**
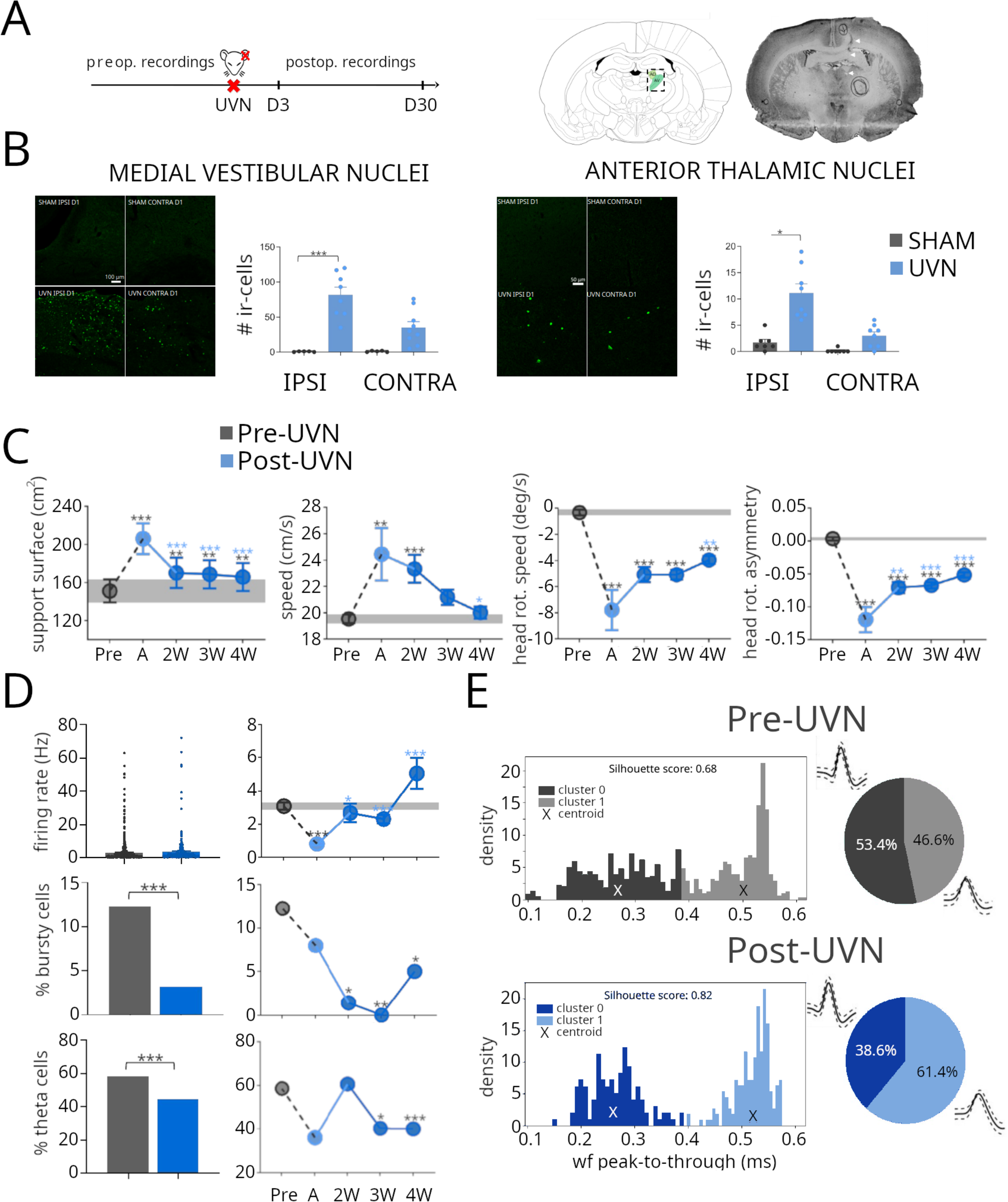
**A**. Left panel: experimental protocol; right panel: coronal section showing the tetrode track in the ATN from one representative animal. **B**. Quantification of post-UVN c-fos immunoreactive cells in the medial vestibular nuclei (MVN, left) and anterior thalamic nuclei (ATN, right) 24h after UVN; left panels: confocal images of MVN and ATN from one sham (top) and one UVN (bottom) rat, the c-fos reactive cells are shown in green; right panels: quantification of the number of c-fos immunoreactive cells in both ipsi-et contra-lesion brain hemispheres. **C**. UVN-induced posturo-locomotor deficits over time (Pre: pre-UVN phase ; A: acute post-UVN phase; 2W-3W-4W: 2, 3 and 4 weeks post-UVN); from left to right: average support surface following the tail-lift test, animal speed, head rotation speed and head rotation asymmetry expressed as the difference between the number of rotations toward the contra- and ipsi-lesion sides divided by the total number of head rotations. **D**. Upper panel: average firing rate; middle panel: percentage of ATN cells firing in bursts; lower panel: percentage of ATN theta modulated neurons; left panels: average values for all neurons recorded during the pre-UVN (grey) and post-UVN (blue) phases; right panels: modifications of the firing properties over time. **E**. Identification of two clusters of ATN neurons based on the waveform characteristics; left panels : distributions of the peak-to-through values for pre-UVN (grey) and post-UVN (blue) neural populations, the silhouette scores of the k-means clustering and the position of the centroids are indicated; right panels: proportion of neurons in the two clusters in pre-UVN and post-UVN neural populations. Black asterisks: statistically significant difference between pre-UVN and post-UVN phases; blue asterisks: statistically significant difference between the acute phase and the subsequent post-UVN phases.

Each recording session lasted 8–10 minutes and was conducted while the animal freely explored the arena. We isolated a total of 849 ATN neurons from five rats preceding the UVN and 308 neurons after UVN (**Fig. 1A**). To ensure that the spike sorting quality was consistent in the pre-and post-lesion periods, we evaluated the isolation quality of the sorted units using the L-ratio, a metric that quantifies how well a cluster (i.e., a putative neuron) is separated from background noise and other units. No significant difference was observed between neurons in the pre-UVN and post-UVN groups (L-ratio: pre-UVN=6.06±0.3, post-UVN=5.86±0.42, unpaired t test with Welch’s correction t=0.37, df=646, adj-p=0.7) showing no significant differences in sorting quality in the pre- and post-UVN periods.

To better characterize the effects of UVN on posturo-locomotor functions and ATN neuronal activity, post-UVN sessions were grouped into four categories according to the time elapsed since the lesion. The first post-UVN week (days 3–6) was defined as the acute phase. Subsequent sessions were grouped into three chronic phases: week 2 (days 7–13), week 3 (days 14–20), and week 4–5 (days 21–33), with week 4 and 5 pooled together due to similar behavioral and neuronal properties. These periods were determined according to the well-characterized time-course of behavioral recovery, reflecting the dynamic of vestibular compensation (Dutheil et al., 2011; El Mahmoudi et al., 2021; Rastoldo et al., 2020).

As previously reported, UVN induced posturo-locomotor deficits that changed across post-lesion phases. First, UVN rats displayed a strong postural instability as attested by an increased support surface, that progressively decreased (**Fig.1C**, first panel from the left; 2-way ANOVA, Animal effect: F(2,8)=254.8, Time effect: F(4,8)=97.32; both p < 0.001) but did not fully recover over time (Tukey’s posthoc test, preop vs acute, vs 1W-2W-3W-4W, adj-p <0.05 for all comparisons). Animal speed was strongly increased after UVN and gradually recovered across post-lesion phases (**Fig.1C**, second panel from the left, 1-way ANOVA, F (4, 668) = 7.116, p<0.001 Tukey’s multiple comparisons test: Pre vs A-2W adj-p<0.01; A vs 4W adj-p<0.05). Finally, rats displayed an increase speed of head rotations toward the ipsi-lesional side (i.e. left side) (**Fig. 1C**, third panel from the left, F (4, 668) = 45.99, p<0.001 Tukey’s test: Pre vs A-2W-3W-4W adj-p <0.001 for all comparisons) as well as an increased number of ipsi-lesional head rotations (attested by head rotation asymmetry, **Fig.1C**, fourth panel from the left, F (4, 668) = 77.25, p<0.00, Tukey’s test: Pre vs A-2W-3W-4W adj-p <0.001 for all comparisons; A vs 2W-3W-4W adj-p <0.01 for all comparisons), that recovered only partly over time. Overall, the UVN-induced behavioral effects are consistent with previous studies and confirm the time-course of posturo-locomotor recovery observed during vestibular compensation (Marouane et al., 2020; Rastoldo et al., 2020).

### Dynamic alterations of ATN firing properties following UVN

We then compared the activity of pre-UVN and post-UVN ATN neurons and observed significant alterations of the basic firing properties following UVN. Although the mean firing rate did not differ significantly between pre- and post-UVN recordings (unpaired t-test with Welch’s correction t = 0.9, df = 450, adj-p > 0.3), a dynamic analysis revealed a transient reduction during the acute phase, followed by a progressive recovery during vestibular compensation (**Fig.1D**, upper panel, Welch’s ANOVA test W(4, 224.3) = 37.17, p < 0.001; Dunnett’s T3 multiple comparisons test: Pre vs A, adj-p < 0.001; Pre vs 2W-3W-4W, adj-p > 0.1 for all comparisons; A vs 2W, adj-p < 0.05; A vs 3W-4W, adj-p < 0.001 for both). This recovery pattern closely mirrors the trajectory described in the vestibular nuclei (VNs), where the acute phase is characterized by a marked reduction in neuronal excitability on the lesioned side, followed by a gradual restoration over time (Ris et al., 1995; Ris and Godaux, 1998; Zennou-Azogui et al., 1994). These results suggest that the ATN, like the VNs, engages in compensatory mechanisms that allow the restoration of neuronal excitability.

To further characterize ATN neuronal activity, we assessed whether cells exhibited burst firing, based on inter-spike interval (ISI) features and burst dynamics. Neurons were classified as bursty according to established criteria for thalamic burst firing in awake animals (Fanselow et al., 2001; Llinás and Steriade, 2006; Ramcharan et al., 2000). Using this approach, we observed a strong and significant decrease in the proportion of bursty cells after UVN, which did not recover over time (Fig. 1D, middle panel; Fisher’s exact test with Benjamini-Hochberg correction: Pre vs A, F = 1.6, adj-p > 0.7; Pre vs 2W, F = 9.8, adj-p < 0.05; Pre vs 3W, F = inf, adj-p < 0.01; Pre vs 4W, F = 2.6, adj-p < 0.05). This persistent reduction in bursty firing likely reflects a disruption in the functional dynamics of ATN neurons following UVN and stands in contrast to the recovered firing rate, suggesting that burst activity is either more sensitive to vestibular input loss or less prone to compensatory plasticity.

A similar trend was observed for the modification of the percentage of theta-modulated cells following UVN. We calculated the intrinsic theta modulation from the autocorrelograms of the firing activity of each neuron. A cell was classified as theta modulated if the power spectrum of the autocorrelogram had a significant peak in the theta band (4-12 Hz). The percentage of theta-modulated cells was significantly and stably reduced after UVN, particularly in late phases (**Fig. 1D** lower panel, Pre vs A, F=2.5 adj-p>0.05; Pre vs 2W, oscillations are generally associated with locomotion, exploration and memory processes (Buzsáki, 2005; Kropff et al., 2021; Sławińska and Kasicki, 1998; Winson, 1978) and previous studies have highlighted a strong link between vestibular information and theta rhythm (Jacob et al., 2014; Russell et al., 2006). The loss of theta-modulated cells following UVN possibly reflects the inability to correctly integrate locomotion and exploration signals, that are necessary for spatial navigation and memory processes. The post-UVN long-lasting impairments of spatial memory that we reported in a recent study support this hypothesis (El Mahmoudi et al., 2023).

### Two electrophysiological populations in the ATN defined by spike waveform

Finally, we analysed whether the modifications of neuron firing patterns may be reflected on the shape of the extracellular action potentials. We quantified for each cell the duration of the average spike, and we extracted two measures: the width of the spike at the half of the peak (half-width, HW) and the distance between the peak and the subsequent trough (peak-to-trough, PT). Neurons recorded during the post-UVN phase exhibited significantly broader spike waveforms compared to pre-UVN neurons, as reflected by increased half-width (HW) and peak-to-trough (PT) durations (HW, pre-UVN = 0.23 ± 0.002; post-UVN = 0.24 ± 0.002, t = 2.6, df = 864, adj-p = 0.01; PT, pre-UVN = 0.37 ± 0.004; post-UVN = 0.42 ± 0.007, t = 5.02, df = 544, p < 0.001). Interestingly, the distribution of PT values displayed a clear bimodal structure both before and after the lesion (**Fig.1E**; Dip’s test: pre, D = 0.03, p < 0.001; post, D = 0.07, p < 0.001), suggesting the coexistence of two distinct neuronal populations—one characterized by short-duration spikes (e.g., dark grey or blue) and the other by longer action potentials (light grey or blue). To objectively validate this separation, we performed K-means clustering (*k = 2*), which confirmed the presence of two clusters with good separation (silhouette score: pre = 0.68; post = 0.82). This supports the existence of two waveform-defined subpopulations within the ATN, potentially corresponding to different functional neuronal types.

Accordingly, the two cell clusters displayed different firing properties. Cells in cluster 0 showed greater firing rate (**Suppl. Fig. 1A**; t=4.78, df=529, adj-p<0.001). In addition, bursty neurons were almost exclusively observed in the first cluster (**Suppl. Fig. 1B**; Chi-2 stat=85.28, p<0.001), whereas the proportion of theta modulated cells was similar between the two clusters (Chi-2 stat=2.49, p=0.1). Cluster 0 also contained a significantly higher proportion of head-direction cells (Chi-2 stat=13.9, p<0.001), position selective cells (Chi-2 contrast, angular head velocity cells were evenly distributed across clusters (Chi-2 stat=1.9, p>0.1) (**Suppl. Fig. 1C**). To further validate the clustering, we trained a random forest classifier using the firing properties and functional cell types described above. The resulting model demonstrated strong performance (**Suppl. Fig. 1D**), further supporting the existence of two distinct subpopulations based on firing properties and functional cell types.

Pre-UVN cells were homogeneously distributed between the two clusters (**Fig.1E**, upper right panel). In contrast, the proportion of post-UVN cells in the first cluster was significantly reduced (**Fig.1E**, lower right panel; Chi-2 stat=19.0: pre-UVN vs post-UVN, p<0.001), suggesting that UVN has a different impact on ATN neuron populations. Altogether, these results indicate that UVN induces long-term functional alterations of neuron firing properties in the ipsilesional ATN. Moreover, they suggest the existence of two distinct cell populations in the ATN that are differently controlled by vestibular input.

### Functional specialization among ATN neurons

We next sought to determine whether UVN could alter the functional properties of the ATN cell population. We identified 4 different cell categories based on classical criteria (see method section): position selective cells (PC), angular head velocity cells (AHVC), speed cells (SC) and finally head-direction cells (HDC). Together, these functionally defined neurons accounted for 52.89% of the ATN population during the pre-UVN phase. We then examined whether these functional categories were differentially affected by UVN.

The proportion of PC was highly similar between the pre- and post-UVN neurons (**Fig. 2A**; Z=0.2, adj-p>0.8), as well as their spatial selectivity (**Fig.2B**, first and second panel from the left; spatial coherence, t=1.4, df=48.4, adj-p>0.1; information content, t=1.3, df=65.1, adj-p>0.1). PC firing activity was not modified (**Fig. 2B**, third panel from the left, t=0.84, df=40.7, adj-p>0.1), but theta rhythmicity was drastically reduced (Z=5.5, adj-p<0.001) similarly to the entire ATN cell population.

**Figure 2:**
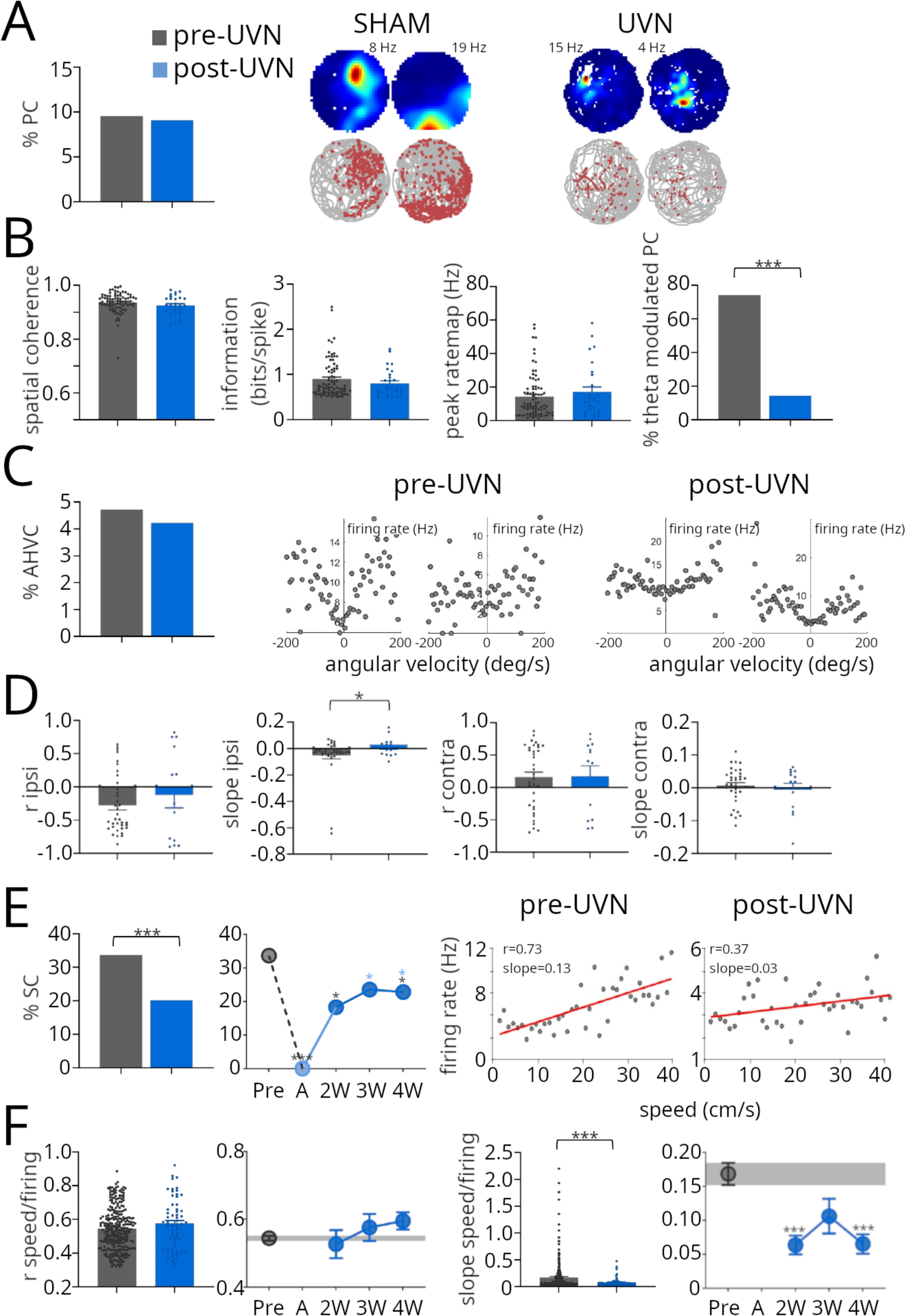
Functional cell categories in the pre-UVN and post-UVN phases. **A**. Left panel: percentage of position-selective cells (PC) in pre-UVN (grey) and post-UVN (blue) neural populations; right panel: 2 examples of PC from one sham and one UVN rat, in the upper panels the color-coded ratemaps show the variation of the firing activity according to the animal position (red indicates highest firing rate) and in the lower panel the trajectories of the rats (in light grey) with the spike position (in red) superposed are shown. **B**. From left to right: average spatial coherence, information content, ratemap peak and percentage of theta-modulated PC, for pre-UVN (grey) and post-UVN (blue) neurons. **C**. Left panel: percentage of angular head velocity cells (AHVC) in pre-UVN (grey) and post-UVN (blue) neurons; right panel: examples of one AHVC from pre-UVN (left) and post-UVN (right) cell populations, the diagrams show the firing rate as a function of angular head velocity bins. **D**. Average correlation value (r) and slope of the regression line between AHVC firing rate and angular head movements toward the ipsi-lesion (first two figures from the left) and contra-lesion sides, for pre-UVN (grey) and post-UVN (blue) neurons. **E**. Left panel: percentage of speed-modulated cells (SC) in pre-UVN and post-UVN neurons (first figure) and variation of SC percentage across pre-UVN and post-UVN phases (second figure); right panel: two examples of pre-UVN and post-UVN SC with diagrams showing the firing rate as a function of animal speed bins (the correlation value and the slope of the regression line are indicated). **F**. From left to right: average correlation value (r) between pre-UVN and post-UVN neuron firing rate and animal speed bins, variations of r across pre-UVN and post-UVN phases, average slope of the regression line for the entire pre-UVN and post-UVN neural populations and across the pre-UVN and the different post-UVN phases. Black asterisks: statistically significant difference between pre-UVN and post-UVN phases; blue asterisks: statistically significant difference between the acute phase and the subsequent post-UVN phases.

AHVC proportion remained constant after UVN (**Fig.2C**; Z=0,3, adj-p>0,7), and maintained the same overall correlation with head-rotation (**Fig.2D**, first and third panels from the left; t=0.7 adj-p>0.4 for ipsi-lesion rotations, t=0.1 adj-p>0.9 for contra-lesion rotations), although the slope of the correlation between their firing and head rotations toward the ipsi-lesion side was significantly altered (**Fig. 2D**, second and fourth panel from the left, t=2, df=45, adj-p=0.05 for ipsi-lesion rotations, t=0.58, df=17.2, adj-p>0.5 for contra-lesion rotations).

The SC was the population mostly affected by the lesion. Indeed, their proportion was significantly and durably lowered during the post-UVN period (**Fig. 2E**; Pre vs A adj-p<0.0001; Pre vs 2W adj-p<0.05; Pre vs 3W adj-p>0.1; Pre vs 4W adj p<0.05) and, although they maintained a similar correlation between their firing rate and animal speed (**Fig. 2F**, first and second panels from the left; t=1.5 adj-p>0.1), the slope was significantly reduced (**Fig. 2F**, third and fourth panels from the left; t=4.7 adj-p<0.001). This result is coherent with the reduced intrinsic theta rhythmicity and reinforce the hypothesis that UVN strongly impacts the integration of locomotor signals within the ATN neural network.

### Directional coding by HDCs is partially restored after UVN

We next focused on HDCs, the most prominently characterized population within the ATN. To overcome the limitations of standard identification criteria following UVN, we implemented a supervised classification approach. HD cells were first selected in the pre-lesion condition using classical criteria (Rayleigh test p < 0.001, vector length > 0.2, and peak firing rate > 1 Hz). These labelled neurons were used to train a random forest classifier based on a set of features capturing directional tuning, burst dynamics, and ISI variability. The resulting model demonstrated excellent performance (**Fig. 3A, right panel**), confirming its reliability for identifying HD cells. This predictive approach, inspired by prior work in sensory-deprived systems (Asumbisa et al., 2022), was then applied to the post-lesion dataset to identify HD cells whose properties were altered and no longer met canonical thresholds.

**Figure 3:**
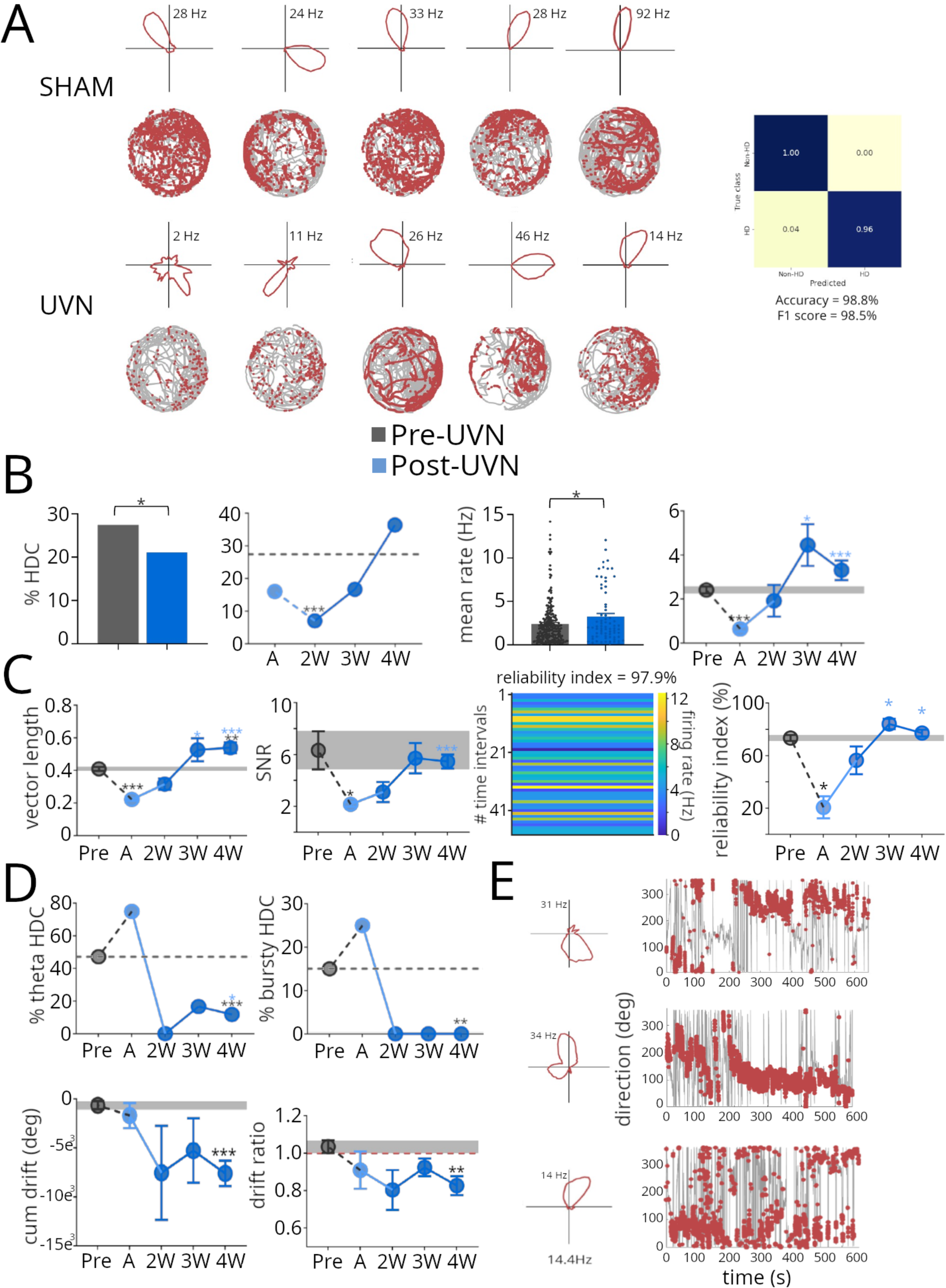
Head-direction cell (HDC) properties in pre-UVN and post-UVN phases. **A**. Examples of HDCs from sham and UVN rats, the upper panel shows the polar plots and the lower panel the trajectory of the rats (in light grey) with spikes (in red) superposed; on the right, the results from the random forest classifier, trained on pre-UVN neurons and used to percentage of HDCs in pre-UVN (grey) and post-UVN (blue) neurons (first figure) and variation of HDC percentage across pre-UVN and post-UVN phases (second figure); right panel: average firing rate of pre-UVN and post-UVN HDCs (third figure) and variations across the different phases (fourth figure). **C**. From left to right: average vector length (first figure), signal to noise ratio (SNR, second figure) and firing reliability index (fourth figure) of pre-UVN and post-UVN HDCs; an example of the plot used to calculate the firing reliability of one HDC is shown (third figure, see methods); the plot shows the color-coded firing rate of the HDC in time intervals (minimum duration 500ms) in which the rat head pointed to the preferred firing direction of the HDC; the firing reliability index is expressed as the percentage of time intervals in which the HDC was active. **D**. Upper panel: percentage of theta-modulated HDCs (left) and HDCs firing in burst (right) across pre-UVN and post-UVN phases; lower panel: average cumulative HDCs drift (left) and average ratio between HDCs drift toward the ipsi-lesion and contra-lesion sides (right) across pre-UVN and post-UVN phases. **E**. Three examples of post-UVN HDCs with polar plots (left) and directions of head movements (in light grey) with spikes (in red) superposed (right). Black asterisks: statistically significant difference between pre-UVN and post-UVN phases ; blue asterisks: statistically significant difference between the acute phase and the subsequent post-UVN phases.

Surprisingly, we observed that HDCs were still present following the lesion (**Fig. 3A**), although their proportion was significantly reduced. This reduction was evident in the early post-lesion phase (i.e. 2W) and gradually recovered over time (**Fig. 3B**, first and second panels from the left; Pre vs A, F=1.9 adj-p>0.4; Pre vs 2W, F=5 adj-p<0.001; Pre vs 3W, F=1.9 adj-p>0.1; Pre vs 4W, F=0.7 adj-p>0.05). This result contrasts with previous studies showing that vestibular impairments abolish the directional firing of HDCs. However, this discrepancy could be attributed to differences in the vestibular lesion models. Typically, vestibular deficits are induced by bilateral lesions of the peripheral vestibular apparatus or by genetic modifications affecting otolith or semicircular canal function (Stackman and Taube, 1998; Valerio and Taube, 2016; Yoder and Taube, 2009), both of which lead to permanent vestibular dysfunction. In contrast, unilateral vestibular loss induces a transient vestibular syndrome, which is followed by central compensation mechanisms involving both The progressive restoration of HDCs we observed during the late post-lesion phases may thus reflect such compensatory plasticity, potentially mediated by a reorganization of sensory inputs to the lesioned ATN — including signals from the intact contralesional vestibular system as well as from non-vestibular sensory modalities. Accordingly, we observed that HDCs firing rate significantly increased when comparing pre- and post-UVN conditions (t = 2.1, df = 98.4, adj-p < 0.05). However, a closer look at the temporal dynamics revealed a sharp reduction during the acute post-lesion phase, followed by a progressive recovery during the later stages of compensation (**Fig. 3B**, panel 3 and 4 from the left; W(4, 19.66) = 25.22, p < 0.0001; Dunnett’s test: Pre vs A, adj-p < 0.0001; Pre vs 2W, adj-p > 0.9; Pre vs 3W, adj-p > 0.3; Pre vs 4W, adj-p > 0.4; A vs 3W, adj-p < 0.05; A vs 4W, adj-p < 0.0001). These results align with the observed restoration of overall ATN activity during vestibular compensation.

A similar dynamic was observed for the directional signal-noise-ratio (SNR) (**Fig. 3C**, panel 2 from the left; W (4, 23.14) = 10.92, p < 0.0001; Dunnett’s test: Pre vs A, adj-p < 0.05; Pre vs 2W-3-W-4W, adj-p > 0.4; A vs 4W, adj-p < 0.0001). This transient drop in SNR suggests that UVN initially degrades the clarity of directional coding, increasing signal noise, which is later restored through compensatory mechanisms.

In line with this degradation and recovery of directional coding, we also observed significant changes in directional tuning strength, as measured by vector length (VL). Post-UVN HDCs exhibited a marked drop in VL during the acute phase, followed by a progressive increase that eventually surpassed pre-UVN levels during the later stages of compensation (**Fig.3C**, first panel from the left; W(4, 21.25) = 57.58, p < 0.0001; Dunnett’s test: Pre vs A, adj-p < 0.0001; Pre vs 2W, adj-p > 0.2; Pre vs 3W, adj-p > 0.6; Pre vs 4W, adj-p < 0.01; A vs 3W, adj-p < 0.05; A vs 4W, adj-p < 0.0001). VL is a classical measure of directional selectivity that reflects how tightly the firing directions of a HDC clusters around its preferred direction, and it is highly sensitive to spike count. The progressive increase in VL observed post-UVN likely reflects the concurrent rise in firing activity, rather than a true enhancement in directional tuning precision. In line with the VL dynamics, the HDC reliability index was initially disrupted following UVN but progressively returned to pre-UVN values (**Fig.3C**, panel 3 and 4 from the left; W (4, 12.77) = 11.08, p < 0.001; Dunnett’s test: Pre vs A, adj-p < 0.05; A vs 3W-4W, adj-p <0.05). The reliability index measures how consistently a HDC is active at its preferred direction. Its progressive increase after UVN further supports the hypothesis of a gradual restoration of the directional coding.

### Persistent impairments in post-UVN HDCs

Despite the recovery of several key features, HDCs exhibited persistent deficits in late post-UVN phases, suggesting that compensatory mechanisms only partially restore their functional integrity. Post-UVN HDCs showed persistent decreased theta modulation (**Fig. 3D** first upper panel; Z=4.9, adj-p<0.001; Pre vs 4W, F=7 adj-p<0.001) and almost complete abolishment of burstiness (**Fig. 3DC** second upper panel; Z=3.2, adj-p<0.01; Pre vs 4W, F=7 adj-p<0.001). From a functional perspective, post-UVN HDCs showed a persistent drift over time (**Fig. 3D** lower panels and **Fig. 3E**, cumulative drift: W (4, 12.65) = 6.26, p < 0.01; Dunnett’s test: Pre vs 4W adj-p<0.001), particularly toward the ipsi-lesional side (drift ratio: W (4, 13.51) = 3.18, p < 0.05; Dunnett’s test: Pre vs 4W adj-p<0.01).

Altogether, these results indicate that although core directional properties of HDCs progressively recover during vestibular compensation, key features such as theta modulation, burstiness, and directional stability remain durably impaired.

### Two HDC subtypes show distinct functional profiles

We next examined whether all HDCs were equally impacted by the lesion. Based on extracellular spike waveform features, we previously identified two distinct neuronal populations within the ATN. Among pre-UVN HDCs, both waveform-defined populations were represented (**Fig. 4A**), but they exhibited markedly different electrophysiological and functional properties. HDCs in cluster 0 (short-duration spikes) displayed higher average and peak firing rates (**Fig. 4B** panel 1 and 2; mean rate: t = 4.8, df = 529, adj-p < 0.001; peak rate: t = 12.3, df = 582, adj-p < 0.001) and stronger directional tuning, as reflected by greater vector lengths (**Fig. 4C**, panel 1; t = 3.8, df = 216, adj-p < 0.001) compared to cluster 1 (long-duration spikes). They also showed higher SNR values (**Fig. 4C**, panel 2; t = 5.5, df = 208, adj-p < 0.001), and were significantly more likely to be bursty (**Fig. 4B**, panel 3Z = 4.4, adj-p < 0.001) and theta-modulated (**Fig. 4B**, panel 4; Z = 2.9, adj-p < 0.01).

**Figure 4:**
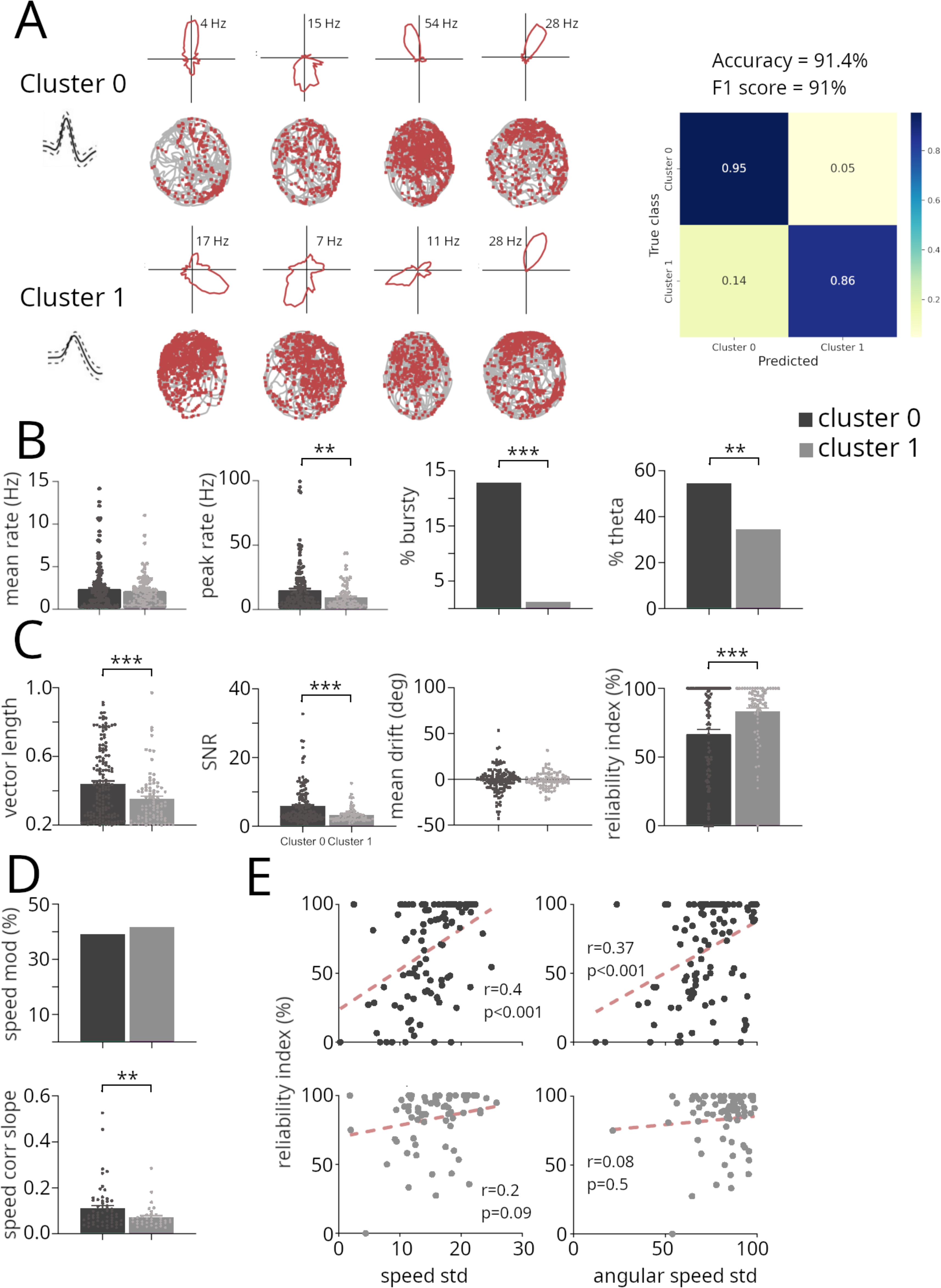
Two cell clusters identified in the pre-UVN HDC population. **A**. Examples of HDCs identified in the two clusters, the upper panel shows the polar plots and the lower panel the trajectory of the rats (in light grey) with spikes (in red) superposed; on the right, the results from the random forest classifier, used to classify pre-UVN HDCs as belonging to either cluster 0 or cluster 1 (see methods). **B**. From left to right: average firing rate, peak rate, percentage of HDCs firing in burst and percentage of theta-modulated HDCs, in the two clusters. **C**. From left to right: average vector length, signal to noise ratio, mean drift and reliability index of HDCs within the two clusters; **D**. Percentage of HDCs modulated by animal speed (upper figure) and average slope of the regression line between firing rate and animal speed, for HDCs within the two clusters; **E**. Scatter plots showing the HDCs reliability index as a function of linear speed variability (expressed as the standard deviation of the speed calculated in a recording session, left panel) or angular speed variability (right panel) for cluster 0 HDCs (upper panel) and cluster 1 HDCs (lower panel); each dot represents one HDC recorded during one training session. Black asterisks: statistically significant difference between cluster 0 and cluster 1 HDCs.

Interestingly, despite this enhanced directional coding, cluster 0 HDCs exhibited lower firing reliability than cluster 1 (**Fig. 4C**, panel 4; t = 3.9, df = 171, adj-p < 0.001), indicating that they were less likely to fire consistently when the animal’s head entered their preferred direction. In addition, they displayed higher speed-tuning slopes (**Fig. 4D**; t = 2.6, df = 89.4, adj-p = 0.01), and their firing reliability was significantly correlated with the variability (standard deviation) of both linear and angular velocity—a relationship absent in cluster 1 (**Fig. 4E**). When exposed to visual cue rotation, cluster 0 neurons also showed significantly higher gain values than those in cluster 1 (cluster 0: 0.75±0.05; cluster 1: 0.46±0.06; t = 3.5, df = 49, adj-p < 0.01), indicating stronger visual anchoring. These findings suggest that cluster 0 HDCs are more strongly driven by visuo-locomotor integration, potentially supporting more flexible but less stable directional representations.

Altogether, these distinct features enabled robust discrimination between the two HDC subtypes. A random forest classifier trained on their electrophysiological and functional signatures achieved strong performance (**Fig. 4A**, right panel), further supporting the existence of two distinct HDC subpopulations within the ATN.

### HDC subtypes are differentially affected by UVN

These functional asymmetries between the two cell clusters raised the possibility that the two HDC subtypes might respond differently to vestibular impairment. We therefore examined whether UVN affected these two populations in distinct ways. Strikingly, we observed opposing dynamics in their relative prevalence: the proportion of HDCs in cluster 0 significantly increased after UVN, while cluster 1 HDCs were strongly reduced (**Fig. 5A, left panel**; cluster 0: Z = 2.02, adj-p < 0.05; cluster 1: Z = –4.18, adj-p < 0.001). A temporal analysis further revealed that cluster 0 neurons remained consistently represented across all post-lesion phases and showed a significant increase during the later stages of compensation (Pre vs acute–2W–3W, adj-p > 0.2; Pre vs 4W, adj-p < 0.001), whereas cluster 1 neurons exhibited a marked and sustained decline (Pre vs 2W–3W, adj-p < 0.01) (**Fig. 5A**, right panel).

**Figure 5:**
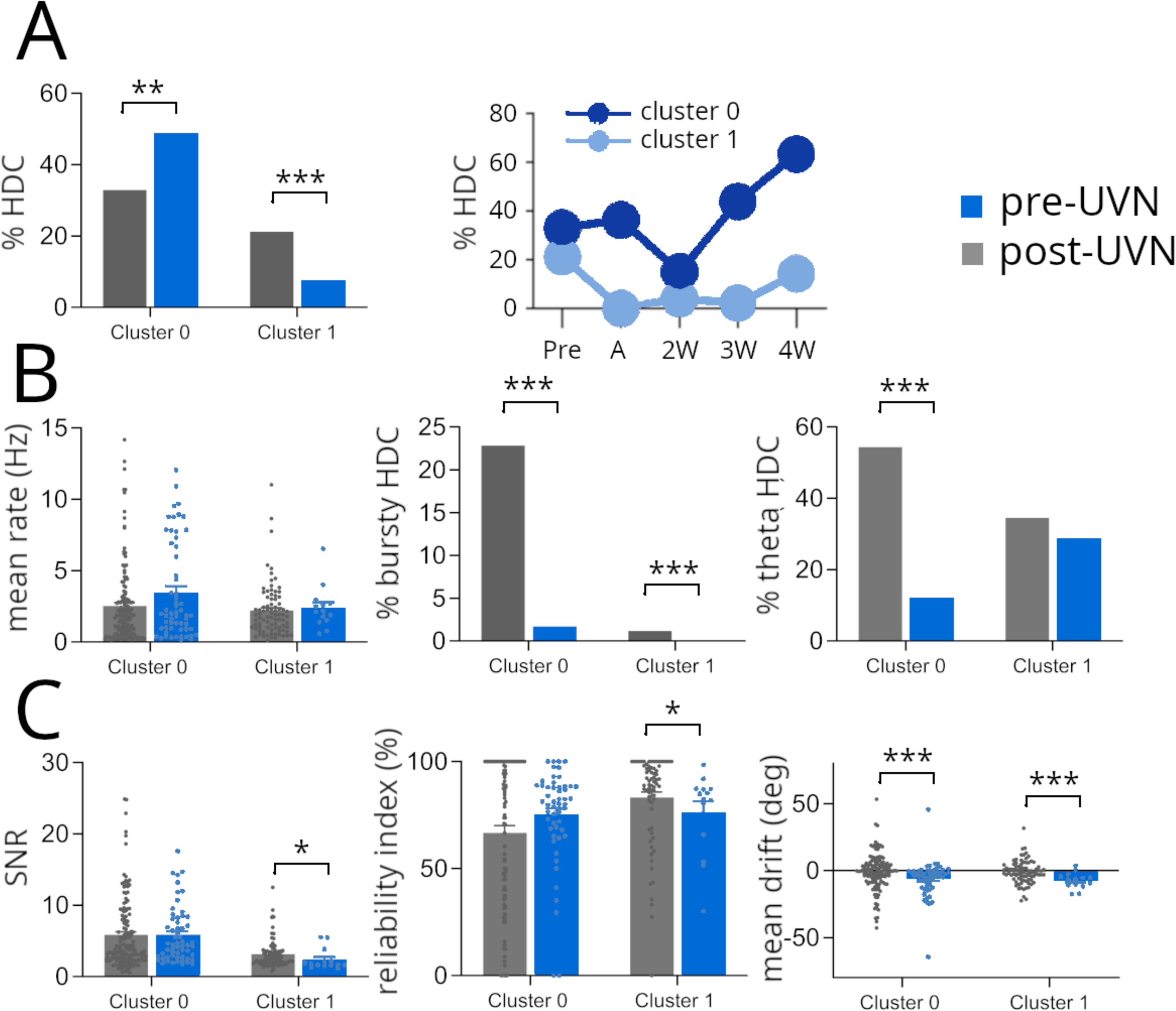
Differences between pre-UVN and post-UVN HDCs within the two clusters. **A.** Left: percentage of pre-UVN (grey) and post-UVN (blue) HDCs identified in the two clusters; right: variations of the percentage of cluster 0 and cluster 1 HDCs across pre-UVN and post-UVN phases. **B**. From left to right: average firing rate, percentage of cells firing in bursts and percentage of theta-modulated cells, for pre-UVN (grey) and post-UVN (blue) HDCs in the two clusters. **C**. From left to right: average signal to noise ratio, reliability index and mean drift, for pre-UVN (grey) and post-UVN (blue) HDCs in the two clusters. Black asterisks: statistically significant difference between pre-UVN and post-UVN phases ; blue asterisks: statistically significant difference between the acute phase and the subsequent post-UVN phases.

We next assessed whether functional properties were differentially impacted by the lesion in the two HDCs clusters. Due to the marked reduction of cluster 1 HDCs during the later post-lesion stages, it was not possible to reliably track functional changes over time within this group. Consequently, all post-UVN phases were pooled for each cluster. While this approach may introduce biases, it allows for a general comparison of pre- and post-lesion functional properties.

First, some similarities were observed between the two clusters. The overall average firing rate of HDCs remained stable in both clusters after the lesion (**Fig. 5B**, first panel; cluster 0: Mann-Whitney U=3862, p>0.2; cluster 1: U=502, p>0.3). Similarly, the proportion of bursty HDCs cells was significantly reduced in both clusters post-UVN (**Fig. 5B**, second panel; cluster 0: Z = -3.6, adj-p < 0.001; cluster 1: Z = –inf, adj-p < 0.001). Both HDCs clusters also showed a strong drift toward the ipsi-lesional side (**Fig. 5C**, third panel, cluster 0: Mann-Whitney U=2183, p<0.001; cluster 1: U=235, p<0.001), reflecting a similar directional instability following UVN.

Despite these similarities, some specific features were differentially impacted by the lesion in the two HDC clusters. The most notable difference was observed in the proportion of theta-modulated cells, that was drastically reduced in cluster 0 whereas remained stable in cluster 1, following UVN (**Fig. 5B**, third panel; cluster 0: Z = -5.5, adj-p < 0.001; cluster 1: Z = –0.4, adj-p > 0.6). In contrast, cluster 1 HDCs showed decreased signal to noise ratio (**Fig. 5C**, first panel; cluster 0: Mann-Whitney U=3862, p>0.1; cluster 1: U=357, p<0.05) and firing reliability (**Fig. 5C**, second panel; cluster 0: Mann-Whitney U=2730, p>0.4; cluster 1: U=299, p<0.05) after the lesion. These results reinforce the hypothesis that the two HDCs clusters represent two distinct neuronal populations that possibly integrate different input types.

Altogether, these findings highlight a differential vulnerability of HDC subtypes to vestibular loss: cluster 1 HDCs were more profoundly affected, showing a marked decline in prevalence and impaired signal fidelity, whereas cluster 0 HDCs appeared more resilient and may contribute to the functional recovery observed during vestibular compensation.

## Discussion

In this study, we investigated how unilateral vestibular neurectomy (UVN) affects neuronal activity in the anterior thalamic nuclei (ATN), with a particular focus on head direction cells (HDCs). We found that UVN induces long-lasting alterations in ATN firing dynamics, including reduced theta modulation, diminished burst firing, and a decreased proportion of speed-modulated cells. Despite these disruptions, HDCs persisted after the lesion and progressively recovered key features of directional tuning—such as firing rate and vector length—during the post-lesion period. However, several core properties, including theta modulation, burstiness, and directional stability, remained impaired even in the late stages of compensation. Moreover, we identified two functionally distinct HDC subtypes that exhibited differential sensitivity to the lesion, with one population being particularly vulnerable to UVN.

While UVN triggered profound and lasting alterations in ATN firing properties, certain features of neuronal activity displayed clear signs of recovery over time. Most notably, we observed a progressive restoration of the mean firing rate within the ipsilesional ATN, which was initially suppressed during the acute post-lesion phase. This rebound mirrors both the progressive recovery of posturo-locomotor functions observed post-UVN, and the recovery of neuronal excitability previously reported in the ipsilesional vestibular nuclei (Ris et al., 1995; Ris and Godaux, 1998), suggesting that shared plasticity mechanisms may operate along the vestibulo-thalamic axis to re-establish baseline firing levels. Consistent with this hypothesis, we observed a strong increase in c-Fos expression within both the medial vestibular nuclei (MVN) and the ipsilesional ATN shortly after the lesion, confirming early and selective engagement of this thalamic region and potentially initiating early compensatory mechanisms. However, this recovery did not extend to temporal features of neural activity: burst firing and theta modulation—both hallmarks of thalamic dynamics— remained durably impaired across all post-lesion phases. This dissociation suggests that, while general excitability can be homeostatically restored, the temporal organization of ATN firing is critically dependent on intact vestibular input. One possibility is that the persistent loss of theta modulation reflects a disruption of hippocampo-thalamic interactions, as ATN theta is known to be modulated by hippocampal outputs via the dorsal fornix (Tsanov et al., 2011). This hypothesis aligns with our recent findings showing long-term impairments in synaptic plasticity within the ipsilesional hippocampus after UVN (El Mahmoudi et al., 2023), suggesting that altered hippocampal output may contribute to the failure of temporal patterning in the ATN.

Interestingly, we were able to identify two distinct electrophysiological populations based on spike waveform, specifically the peak-to-trough duration. This bimodal distribution recapitulates prior observations of ATN neuronal heterogeneity (Laurens et al., 2019) and suggests the existence of different neuronal subtypes within this region, with markedly different electrophysiological and functional profiles: one group (cluster 0) was characterized by shorter spike durations, higher firing rates, greater burst and theta modulation, and stronger tuning to position, direction and speed; the other (cluster 1) had broader spikes and more stable but less selective firing. Importantly, both clusters were consistently observed across the entire dorsoventral extent of the recording tracks, suggesting that this distinction is not due to anatomical sampling differences, but rather reflects intrinsic heterogeneity within the ATN itself. Strikingly, the two populations were differentially affected by UVN. Cluster 0 neurons were the most affected, suggesting that they are reliant on vestibular input. This result raises important questions about the respective roles of the two clusters and their afferent connectivity. One possibility is that these subtypes reflect functionally segregated cell clusters within the ATN, each receiving different patterns of input from upstream and downstream structures. Although speculative, this interpretation aligns with previous work showing that subregions of the ATN receive distinct combinations of sensory and cortical afferents (Jankowski et al., 2013). Such anatomical specialization could help explain the asymmetric vulnerability of these two neuronal populations to vestibular loss.

Our results revealed that the vestibular loss did not equally affect the distinct classes of spatially tuned neurons that have been consistently described in the ATN, namely position-, angular head velocity–, speed-, and head direction-selective cells (Laurens et al., 2019; Lomi et al., 2023; Taube, 1995; Yoder et al., 2011). Among the four functional classes, speed-modulated cells (SCs) were the most profoundly impacted by the lesion. Their proportion dropped significantly following UVN and remained low throughout the compensation period, suggesting a long-lasting disruption in the integration of locomotor signals within the ATN. This finding aligns with the observed reduction in theta modulation, as speed coding and theta rhythmicity are often tightly linked, particularly in thalamo-hippocampal circuits (Hinman et al., 2011; Kennedy et al., 2022). In contrast, position-selective cells (PCs) were largely preserved in terms of spatial tuning and firing rate, although they exhibited a drastic reduction in theta rhythmicity. Similarly, angular head velocity (AHV) cells were stable in proportion but showed a specific and durable alterations in their tuning to ipsilesional head rotations. This physiological asymmetry mirrors the behavioral phenotype observed post-UVN, which included a persistent increase in both the angular speed and frequency of ipsilesional head turns. Unlike postural or locomotor deficits, these AHV-related impairments did not fully resolve over time, suggesting that angular velocity coding is particularly sensitive to unilateral vestibular disruption and may remain chronically imbalanced.

Finally, head direction cells (HDCs) showed a dynamic response to vestibular loss, with a marked reduction following UVN and a progressive recovery over the course of compensation, eventually reaching levels comparable to the pre-lesion condition. This suggests that directional tuning can be re-established through central plasticity mechanisms, potentially involving inputs from intact contralesional vestibular structures or alternative sensory pathways. Nevertheless, the functional restoration over time was not complete since HDCs continued to exhibit persistent reduced theta modulation and burst firing, as well as an increased directional drift. Interestingly, HDCs were observed in the two cell clusters but the effects of UVN highly diverged between the two. Prior to the lesion, the two HDC populations exhibited distinct functional profiles: cluster 0 HDCs showed stronger directional tuning, greater theta modulation, and more sensitivity to linear and angular velocity, compared to cluster 1 HDCs. In addition, cluster 0 neurons exhibited greater sensitivity to visual cue rotations, suggesting a tighter coupling to visuo-locomotor dynamics. Following UVN, cluster 1 HDCs showed a sharp and sustained decrease in prevalence and tuning quality, while cluster 0 HDCs were more resilient and contributed disproportionately to the recovery of directional coding over time. This differential vulnerability suggests that vestibular input is not equally critical for all HDCs, and that these subtypes may support complementary facets of directional processing—some more dependent on flexible, multimodal integration, others more anchored to vestibular input. Notably, cluster 0 HDCs not only persisted but exhibited changes in their tuning properties following the lesion, suggesting they may undergo plastic modifications to compensate for the loss of vestibular input. This adaptation could involve increased reliance on non-vestibular signals, such as proprioceptive, visual, or motor efference cues, as well as residual input from the contralesional vestibular nuclei. The fact that unilateral, but not bilateral, vestibular lesions allow for partial recovery of directional coding further supports the idea that the intact contralateral vestibular system likely plays a key role in this process. To our knowledge, this is the first demonstration that distinct subpopulations of HDCs within the ATN exhibit differential vulnerability to vestibular loss, unveiling a new layer of functional diversity in the head direction system.

## Conclusion

Together, our findings shed new light on the plasticity of the head direction system in response to unilateral vestibular loss. By combining histological, behavioral, electrophysiological, and machine-learning approaches, we reveal a complex reorganization of ATN circuits, marked by persistent disruptions in temporal dynamics and selective impairments in spatial coding. The identification of two functionally distinct HDC subtypes, each with a different degree of vulnerability, highlights the heterogeneous dependence of directional neurons on vestibular input. While some HDCs appear capable of adapting to altered sensory environments, others remain critically reliant on intact vestibular signals. These insights advance our understanding of directional coding plasticity and indicate that compensation following vestibular loss is partial and uneven. Future studies will be needed to identify the circuit-level mechanisms underlying this selective reorganization, and to determine whether similar patterns exist in other nodes of the head direction network.

Crucially, our results provide the first direct neurophysiological evidence of how unilateral vestibular loss alters HDC activity, offering a mechanistic framework to understand how directional circuits adapt to asymmetric sensory deprivation. Moreover, they provide a neural basis for the spatial navigation and memory impairments commonly observed in patients with vestibular dysfunction.

## Author contributions

N.E.M., F.S. and P.Y.J. designed the study. P.Y.J and N.E.M performed surgeries. N.E.M performed experimental acquisitions. N.E.M. and A.H. analysed data. J.L provided help in analysing the data. C.L provided experimental and technical support. N.E.M and F.S wrote the manuscript. All authors commented and edited the manuscript.

## Data availability

All data and codes for the analysis are available upon request to the corresponding author

## Acknowledgment

We thank David Péricat for his help with the surgeries. We thank the caretakers of the CRPN for taking care of the animals. We thank the BNCS team for discussions. Financial support for this work was provided by the *Centre National de la Recherche Scientifique* and *Ministère de la Recherche*.

## Methods

### Animals

The study was carried out within the terms of the directive 2010/63/EU of the European Parliament and of the Council, French institutional guidelines AGRG1238767A, and approved by the Ethical Committee 71 (APAFIS #33350). Four Long-Evans male rats (300– 350 g) were used for post-lesion c-Fos immunohistochemistry. Two rats underwent left unilateral vestibular neurectomy (UVN), and two received sham surgery. Animals were perfused 24 hours after surgery for tissue processing. For electrophysiological recordings and behavioral assessments, five Long-Evans male rats (Charles River) weighing 300-350g were housed in individual cages (40 cm long x 26 cm wide x 16 cm high) with ad lib food and water and kept in a temperature-controlled room (20 ± 2 °C) with natural light/dark cycle. One week after arrival, animals were handled daily by the experimenter for 7 days. All rats received surgery for tetrodes implantation into the left anterior thalamus nuclei (ATN). Following 7 days of recovery, behavioral trainings and ATN’s activity recordings started. Rats were placed on a food-deprivation schedule to keep them at 90% of their free-feeding body weight. ATN’s recordings were performed for 4 pre-lesion weeks. At the end of this period, food-deprivation was stopped, and rats were submitted to left unilateral vestibular neurectomy (UVN). Following 2-3 days of recovery, training started again until the 5^th^ post-lesion week (33 days following UVN).

### Surgery

#### Microdrives and tetrodes implantation

Recordings were made using tetrodes, each composed of four twisted 25µm or 17µm polyumid-coated platinum-iridium (90%/10%) wires (California Fine Wire, CA), attached to an Axona 32 channels microdrive (Axona Ltd, Herts, UK). A subcutaneous injection of buprenorphine (Buprecare®; 0.02 mg/kg) was realized 30 minutes before the beginning of the surgery. Anesthesia was induced with isoflurane (3%-3.5%). Tetrodes were chronically implanted in the left ATN (AP: −1.8, ML: ± 1.4, DV: −3.9). Before awakening, animals were injected subcutaneously with a solution of Ringer Lactate (Virbac; 10 ml/kg) to alleviate the dehydration resulting from the surgery and with antibiotic (oxytetracycline, 10mg/kg) to prevent from infections.

#### Unilateral vestibular neurectomy (UVN)

Unilateral vestibular loss was induced by left UVN consisting in sectioning the left vestibular nerve. Briefly, animals were anesthetized with isoflurane (4%) 30 minutes after a subcutaneous injection of buprenorphine (Buprecare®; 0.02 mg/kg). The animals were intubated, and the anaesthesia was maintained during the surgery with isoflurane (3%). To access the left vestibular nerve, the tympanic bulla approach was used (see Péricat et al., 2017 for details). The left vestibular nerve was then sectioned at a post-ganglion level, close to the brainstem. Before awakening, animals were injected subcutaneously with a solution of Ringer Lactate (Virbac; 10 ml/kg) to alleviate the dehydration resulting from the surgery. The success of the UVN was attested by the immediate appearance of a characteristic vestibular syndrome composed of postural, locomotor, and oculomotor deficits (Péricat et al., 2017).

### Behavioral and recording experiments

#### Apparatus & experimental protocol

The apparatus was a circular black open field (100 cm diameter, wall 50 cm high) with a plastic floor that was wiped with alcohol before each session to prevent uncontrolled odor cues, and isolated from the rest of the room by an opaque circular curtain. The open field was polarized with a white card (20cm x30cm) attached to the wall. A food dispenser was located 2m above the open field and dropped 20mg of food pellets each time it was activated. A radio, fixed to the ceiling above the open field was used to mask uncontrolled directional sounds. The unit recording system and the experimenter were in an adjacent room. Each recording session consisted of random foraging and was conducted for 8-10 minutes.

#### Behavioural assessments

To assess vestibular function, we evaluated postural stability at different time points after UVN (acute phase – day 3; 2 week – day 7; 3 weeks – day 14; 3 weeks – day 21; 4 weeks – day 30). To do so, we measured the support surface (i.e., the area between the four paws of the animal) after the tail-lift reflex test (TLR) allowing to assess the effectiveness of the vestibulo-spinal reflex (Cassel et al., 2018; Grosch et al., 2021; Martins-Lopes et al., 2019; Tighilet et al., 2017). The support surface was calculated in cm^2^, using an image analysis system developed on Matlab®. For each animal, ten measurements were taken and averaged at different timepoints of the pre-lesion and post-lesion periods.

Animal trajectory and head direction were sampled at 25Hz using two diodes attached to the headstage and stored as x/y coordinates. The total distance travelled, and the average speed were calculated for each session. Head rotations and angular speed toward the ipsi- and contra-lesion sides (i.e. left and right respectively) were also quantified.

#### Neural recordings

Screening and recordings were performed with a counterbalanced cable attached at one end to a commutator, allowing the rat to move freely. The other end of the cable was connected to the headstage, which contained a field effect transistor amplifier for each wire. The signals from each tetrode wire were further amplified 10,000 times, bandpass filtered between 0.3 and 6 kHz with Neuralynx amplifiers (Neuralynx, Bozeman, MT, USA), digitized (32 kHz), and stored by DataWave Sciworks acquisition system (DataWave Technologies, Longmont, CO, USA). Two light-emitting diodes (LED), one red and one green, separated by 5 cm and attached to the headstage assembly provided the position and the orientation of the rat’s head. The LEDs were imaged with a CCD camera fixed to the ceiling above the maze, and their position was tracked at 25 Hz with a digital spot-follower.

### Data analyses

All data analyses (except for the spike sorting) were performed with home-made scripts on Matlab® and Python.

#### Spike sorting

Spike sorting was performed manually using the graphical cluster-cutting software Offline Sorter (Plexon). Units selected for analysis had to be well-discriminated cluster with spiking activity clearly dissociated from background noise. Units having inter-spike intervals < 2 ms (refractory period) were removed due to poor isolation. To prevent repeated recordings of the same cell over days, clusters that recurred on the same tetrodes in the same cluster space across recording sessions were only analysed on the first day. To asses cluster isolation quality we calculated the L-ratio (Schmitzer-Torbert et al., 2005), and we compared these values between pre-UVN and post-UVN neurons using an unpaired t-test with Welch’s

#### Overall ATN neuron firing properties

For each neuron we calculated the **mean firing rate** (total number of spikes divided by the time length of the recording session). The **peak firing rate** was determined as the maximum firing rate observed across 200 ms non-overlapping time bins over the entire session. To assess firing rate stability, the **coefficient of variation (CV) of peak** amplitudes across spikes was also computed.

To assess **burstiness**, we computed inter-spike intervals (ISI) for each neuron and extracted several parameters reflecting the temporal structure of spiking activity. A burst was defined as a sequence of at least two spikes separated by ISIs shorter than 6 ms. For each cell, we calculated the number of bursts, burst rate (bursts per second), mean burst length (number of spikes per burst), and mean intra-burst frequency. A cell was classified as bursty if it met all the following criteria: more than 10% of ISIs shorter than 6 ms, burst rate exceeding 0.05 bursts/s, mean burst length ≥ 2 spikes, and mean intra-burst frequency above 100 Hz. Neurons that did not meet these conditions were further categorized as tonic (regular firing), sparse firing (long ISIs), or mixed firing, based on the relative proportions and variability of their ISI distributions. These thresholds were chosen to reflect typical burst firing patterns observed in thalamic neurons (Fanselow et al., 2001; Llinás and Steriade, 2006; Ramcharan et al., 2000).

The intrinsic **theta modulation** (4-12 Hz) was assessed by calculating the power spectrum of the spike time autocorrelogram (1000ms time window, bins of 10ms) using the fast-Fourier transform. A cell was considered as theta modulated if the power spectrum in the theta band (4-12 Hz) exceeds five times the average value of the entire power spectrum.

We also measured the average waveform of the spikes for each cell cluster, and we extracted two measures: the **width at half of the peak** (HW) and the distance between the **peak and the first subsequent through** (PT).

We assessed the modality of the peak-to-trough (PT) duration distribution using Hartigan’s dip test, to determine whether distinct neuronal subtypes were present. A k-means clustering analysis was then applied to PT values within the pre-UVN and post-UVN populations. The silhouette score was computed to evaluate the quality and consistency of the clustering. The proportion of neurons assigned to each cluster was compared between pre and post lesion periods using a chi-squared test.

To further validate the clustering based on peak-to-trough duration, we trained a random forest classifier using a combination of waveform features (excluding PT) and firing pattern metrics. The model was trained using 250 estimators and balanced class weights to account for potential imbalance between groups. Model performance was evaluated via stratified 10-fold cross-validation, accuracy score, F1-score, and confusion matrix. This approach confirmed the functional relevance of the clustering and identified the most informative features contributing to the classification of neuronal subtypes.

#### Functional classification of ATN neurons

To characterize the functional diversity of ATN neurons, we classified each cell based on its tuning to spatial position, linear speed, angular head velocity (AHV), and head direction (HD), using established criteria from the literature (Góis and Tort, 2018; Graham et al., 2023; Lomi et al., 2023) combined with shuffle-based significance testing.

Position cells were identified by constructing spatial rate maps (in Hz) normalized by the occupancy time of each spatial bin. Only bins visited by the animal were included. Spatial information content was quantified using Skaggs’ information measure (in bits/spike), and spatial coherence was computed using two complementary methods: a direct spatial autocorrelation and a matrix-based approach using the rate and occupancy maps. A neuron was classified as a position cell if it had a peak firing rate > 1 Hz, spatial coherence > 0.6 by both methods, Skaggs information > 0.5 bits/spike, and at least one clearly defined firing field detected as a contiguous region of elevated firing in the ratemap.

Speed cells were identified by examining the relationship between firing rate and the animal’s linear speed, binned in 1 cm/s intervals. For each neuron, we computed the Pearson correlation coefficient between average firing rate and speed and fitted a linear regression model. The analysis was restricted to speeds between 1 and 40 cm/s. Statistical significance was assessed using a shuffle procedure (1000 iterations) in which firing rates were randomly permuted across speed bins. A neuron was classified as a speed cell if it showed a correlation coefficient > 0.3, a regression slope > 0.025, and both the regression and shuffle-based p-values < 0.05.

Angular velocity cells (AHV cells) were identified by computing firing rate as a function of angular head velocity, estimated from the smoothed derivative of head direction. Angular velocity values were binned from −200 to +200 deg/s in 6 deg/s steps. We separately analyzed clockwise (CW) and counterclockwise (CCW) directions, computing the correlation and slope of the firing rate with respect to AHV in each. A neuron was classified as AHV-sensitive if it showed a correlation > 0.3, a slope > 0.025, and exceeded the 95th percentile of a shuffle distribution (1000 permutations) for either direction. AHV cells were further categorized as symmetric, asymmetric, or non-reactive, based on the relative balance of CW and CCW tuning.

Head direction cells were identified by computing directional tuning curves in 6° bins. Directional selectivity was quantified using the mean resultant vector length (Rayleigh vector), and statistical significance was evaluated using both the Rayleigh test and a shuffle-based threshold (1,000 random circular shifts). A neuron was considered HD-tuned if it had a vector length > 0.2, a p-value < 0.001, and a peak firing rate > 1 Hz.

#### HDC activity

The rat’s head direction was computed at each time point from the relative position of two tracking LEDs projected onto the horizontal plane. For each neuron, a directional tuning curve was generated by dividing the number of spikes emitted when the animal was facing a given direction (in 6° bins) by the time spent facing that direction.

The preferred firing direction (PFD) was defined as the angle corresponding to the bin with the highest firing rate. The peak firing rate was calculated as the maximum of this tuning curve, and the mean firing rate as the average across all direction bins. The strength of directional tuning was quantified using the Rayleigh vector length, and the tuning width was estimated by fitting a centered Gaussian to the tuning curve and calculating the full width at half maximum (FWHM) of the fit. The signal-to-noise ratio (SNR) was defined as the ratio between the mean firing rate inside the tuning range and the mean rate outside of it.

To assess the temporal stability of directional tuning, we computed a reliability index based on the proportion of time intervals during which the animal was oriented within the Gaussian-defined tuning range and the neuron was active. To quantify potential drift in directional tuning, we computed the difference in PFD between spikes separated by fixed intervals of 100 spikes. From this, we extracted the mean angular drift between spikes and the total cumulative drift across the session. We also quantified total angular deviations to the left and right and computed a contra-/ipsilateral deviation ratio to evaluate any directional bias in drift.

When an HDC was identified, we performed a second recording session with a rotated visual cue and calculated the HDC activity gain by measuring the shift in the neuron’s PFD following the cue rotation. The gain was defined as the absolute ratio between the angular shift in the PFD and the imposed rotation angle of the cue (Asumbisa et al., 2022). This metric allowed us to quantify the extent to which the neuron’s directional coding followed the external cue, with a gain of 1 indicating perfect cue anchoring and a gain of 0 indicating no influence of the cue on the cell’s tuning.

#### Validation of pre-lesion HD cell subtypes

To assess whether the two identified subpopulations of HDCs could be distinguished based on their functional and electrophysiological features, we trained a supervised classification model on the pre-lesion dataset. We selected only neurons that were classified as HDCs (Rayleigh vector length > 0.2, peak firing rate > 1 Hz), and trained a Random Forest classifier using 12 features spanning multiple domains: directional tuning (Rayleigh vector length, peak and mean HD firing rates, Gaussian tuning width, SNR), theta modulation, firing pattern (burstiness, intra-burst frequency, ISI variability), speed modulation, and reliability of firing within the tuning range. The model was trained using 250 estimators, class balancing, and square-root feature sampling at each split. Model performance was evaluated via 10-fold stratified cross-validation. Classification accuracy, F1-score, and confusion matrix were computed to assess the model’s generalization performance. Feature importance was extracted from the trained classifier to identify the most discriminative variables separating the two HDC clusters.

#### Post-lesion HD classification

To identify HDCs in the post-lesion period, we trained a supervised machine learning model on the pre-lesion dataset. A random forest classifier was fitted using 13 functional and autocorrelogram-derived features, including directional tuning strength (e.g., Rayleigh vector length, peak HD rate, mean HD rate), bursting dynamics (e.g., mean burst length, burst rate, intra-burst frequency), firing variability (e.g., CV of ISI), and temporal structure metrics derived from the spike train autocorrelogram (e.g., peak amplitude, peak frequency). The classifier was trained using 250 estimators with square-root feature sampling at each split. Model performance was evaluated using stratified 10-fold cross-validation, accuracy, F1-score, and confusion matrix. Once validated, the trained model was applied to the post-lesion dataset to predict HDC identity, providing an independent, feature-based estimate of directional tuning in the absence of direct Rayleigh-based labels.

### Post-recordings histology

Animals were sacrificed at the end of the experiments for histological investigations by injections of Euthasol® (300mg/kg, *ip*). The brains were removed and frozen with dry ice. 40µm-thick coronal sections were mounted on glass slides and stained with thionine. The position of the tips of the electrodes was determined from digital pictures, acquired with a Leica Microscope (Wetzlar, Germany).

### C-fos immunohistochemistry

Immunohistochemical labelling was performed according to previously validated protocols (Dutheil et al., 2016; El Mahmoudi et al., 2022, 2021; Tighilet et al., 2007). Briefly, rats were perfused transcardially 24 hours after surgery (UVN or sham) with 0.9% saline followed by 4% paraformaldehyde (PFA) in 0.1 M phosphate buffer (PB). Brains were post-fixed for 24 hours and cryoprotected in 30% sucrose solution at 4°C. Coronal sections (40 μm) were cut using a freezing microtome and stored in cryoprotectant at –20°C until processing. Free-floating sections were rinsed in 0.1 M phosphate-buffered saline (PBS, 3×5 min), then saturated and permeabilized in a solution containing 5% bovine serum albumin (BSA) and 0.3% Triton X-100 for 1 hour. Sections were incubated for 24 hours at 4°C with a rabbit anti-c-Fos primary antibody (1:5000, Sigma-Aldrich, Ab-5) diluted in PBS containing 5% BSA and 0.3% Triton X-100. After rinsing in PBS, sections were incubated with a goat anti-rabbit secondary antibody conjugated to Alexa Fluor 488 (1:500; Invitrogen, A11008) for 1 hour at room temperature. Sections were mounted with Roti®-Mount FluorCare medium (Roth, HP19.1). Confocal imaging was performed using a Zeiss LM 710 NLO laser scanning microscope equipped with a 20X or 40X objective. c-Fos-positive nuclei were manually counted in the medial vestibular nuclei (MVN) and anterior thalamic nuclei (ATN) on 3–5 sections per animal, with anatomical localization verified using a rat brain atlas (Paxinos and Watson, 2009). Cell counts were averaged per region for statistical analysis.

### Statistical analysis

Immunohistochemistry data were analysed using a non-parametric Kruskal–Wallis test to compare the number of c-Fos positive cells across groups (sham and UVN, both ipsilesional and contralesional sides), followed by Dunn’s multiple comparisons test when appropriate. Behavioral parameters—including support surface, average speed, head rotation speed, and rotation asymmetry (ipsi-vs. contra-lesion)—were compared between pre- and post-UVN periods using a two-way ANOVA for support surface, and one-way ANOVAs followed by Tukey’s post hoc tests for the other variables. Neuronal data were analysed using unpaired t-tests with Welch’s correction for pre-vs. post-lesion comparisons. For analyses involving HDC subtypes (clusters 0 and 1), non-parametric Mann–Whitney U tests were used. Temporal dynamics across post-lesion timepoints were assessed using Welch’s ANOVA followed by Dunnett’s T3 multiple comparisons test. The proportions of functionally classified cells were compared using Z-tests for proportions (with Benjamini–Hochberg correction for multiple comparisons), and Fisher’s exact tests were used for comparisons across pre- and post-lesion phases. All results were considered statistically significant at p < 0.05.

---

**Supplementary figure 1: A**. Average firing rate (left), peak firing rate (middle) and coefficient variation of the peak firing (right) for all pre-UVN ATN neurons in the two clusters. **B**. Percentage of theta-modulated cells and cells firing in bursts for pre-UVN neurons in the two clusters. **C**. Percentage of head-direction cells (HDC), position-selective cells (PC), angular head-velocity cells (AHV) and speed cells (SC) identified in the pre-UVN ATN cell population and belonging to the two clusters. **D**. Results from the random forest classifier, used to classify pre-UVN ATN cells as belonging to either cluster 0 or cluster 1 (see methods). Black asterisks: statistically significant difference between cluster 0 and cluster 1 neurons.

