## Supplementary figures and images for "Dynamics of thalamic directional coding under vestibular imbalance"

### Supplementary figure 1

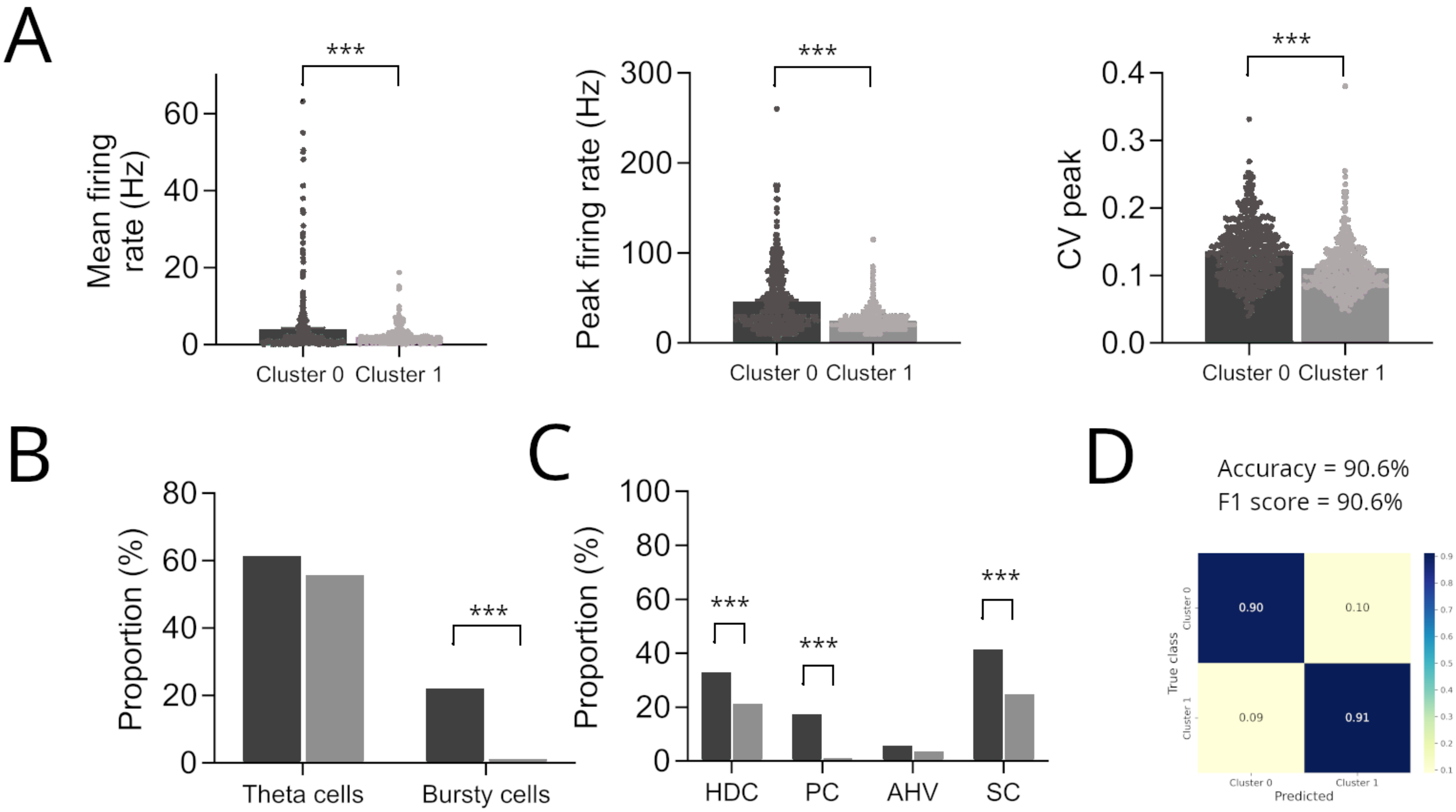
